# Metabolic and genomic adaptations of *Salmonella* Typhimurium grown on itaconate

**DOI:** 10.1101/2025.06.20.660670

**Authors:** Jacob Pierscianowski, Suzana Diaconescu-Grabari, Farhan R. Chowdhury, Andréanne Lupien, Brandon L. Findlay, Karine Auclair

## Abstract

*Salmonella enterica* ser. Typhimurium (STm) is an enteric pathogen that causes almost 100 million cases of salmonellosis a year. A hallmark of STm is their ability to survive and replicate in the macrophages that phagocytose them. Inside the *Salmonella*-containing vesicles (SCV) STm is exposed to nutrient limitation and antimicrobial molecules, including itaconate, a dicarboxylate produced at millimolar concentrations by activated macrophages. STm has an itaconate degradation pathway that allows it to detoxify itaconate and convert it into pyruvate and acetyl-CoA. In the SCV, STm undergoes extensive metabolic reprogramming in response to the nutrient limitation and high stress environment, and strict catabolite control is employed at various stages of infection, depending on carbon source availability. The extent to which itaconate is used as a carbon source by STm is not currently well understood. Here, we explore the ability of STm to grow in media containing itaconate as the sole carbon source, identify the genes of the itaconate response operon necessary for itaconate degradation, and confirm their importance during infection of macrophages. Growth on itaconate was substantially slower than in carbon-rich or acetate media, with multiple subcultures allowing for increased growth rates. Genomic analysis suggests that mutations in the RpoS-stress response and increased expression of the DctA transporter improve growth on itaconate. Using metabolomics, it was revealed that growth on itaconate resulted in substantially reduced levels of gluconeogenesis compared to growth on acetate and increased glutathione disulfide levels, indicating a high level of oxidative stress. Overall, our results point towards the STm itaconate degradation pathway having a more important role in detoxifying itaconate, rather than using it as a carbon source inside the macrophage. Additionally, this study establishes media conditions that would enable the high throughput discovery of potential inhibitors of itaconate degradation.

## Introduction

*Salmonella enterica* serovar Typhimurium (STm) is a facultative intracellular pathogen that causes gastroenteritis in humans, with the potential for systemic infection. It is the primary causative agent of non-typhoidal *Salmonella* infections, with over 93 million cases and over 155 000 deaths yearly worldwide, earning it a classification as a high priority pathogen by the WHO [1–4]. During infection, STm survives in a diverse range of environments, including inside host macrophages, in specialized compartments called *Salmonella*-containing vesicles (SCV) [5–7]. The SCV is a nutrient-limited, mildly acidic (pH 4-5), and oxidative environment that STm has evolved to survive in, thanks in part to its metabolic flexibility allowing the use of a variety of carbon sources [8]. While many studies have added to our understanding of which metabolic pathways are used by intracellular STm both *in vivo* and *in vitro*, the full picture remains incomplete [5–7]. To some degree, this is due to the heterogeneity of STm physiology during infection, even when infecting the same tissue or cell type [9,10]. This localization of bacteria during infection is of particular importance because it offers a protective environment from some antibiotic treatments; owing to both limited accessibility by antibiotics with poor membrane-permeability and the presence of slow growing STm populations that are resistant to treatment [10,11]. Nutrient starvation has been shown to be the primary cause of this protective phenotype, even when compared to other stresses, such as acidic pH or oxidative stress, and can be attributed to the entire population, not just a small subsect of highly-resistant persisters [10,12,13]. As such, recognizing how, when, and why STm remodels its metabolism in response to nutrient limitation, will afford a better understanding of why antibiotic treatments might fail and how to design new drugs targeting intracellular infections [14].

Itaconate is an antibacterial and immunomodulatory molecule produced by activated macrophages and pumped into the SCV at concentrations as high as 8 mM [15,16]. The antimicrobial activity of itaconate is diverse; it can act as a non-covalent inhibitor, as well as a covalent modifier of cysteine residues in proteins through “*S*-itaconations”, which can result in the inhibition of enzyme activity or modulation of protein function [17,18]. Over 1000 proteins in STm alone have been shown to be subject to *S*-itaconation, including enzymes involved in the TCA cycle (SDH: succinate dehydrogenase, SCS: succinyl-CoA synthetase) [19,20], in the glyoxylate shunt (Icl: isocitrate lyase) [21–23], and in *de novo* purine biosynthesis (PurF: amidophosphoribosyltransferase, GuaA: GMP synthetase, GuaB: IMP dehydrogenase, GuaC: GMP dehydrogenase) [24,25]. In response, STm and several other intracellular pathogens including *Pseudomonas aeruginosa* and *Mycobacterium tuberculosis* (Mtb), have evolved enzymes for itaconate detoxification [25–27]. The itaconate degradation pathway consists of three enzymes that convert itaconate into acetyl-CoA and pyruvate (**Fig 1B**): itaconate CoA transferase (Ict), itaconyl-CoA hydratase (Ich), and citramalyl-CoA lyase (Ccl) [26]. The first step, involving the formation of itaconyl-CoA, has also been found to be catalyzed by other transferases or by SCS [28–31]. In STm, the three genes coding for the itaconate degradation enzymes (**Fig 1A**) are found within the same operon along with one other gene, *lgl*, which encodes a lactoylglutathione lyase involved in the detoxification of methylglyoxal [32–34]. Lgl is a secondary glyoxalase I enzyme, performing the same function as the canonical homolog in STm, GLO1, which is part of the methylglyoxal pathway along with GLO2 (**Fig 1C**). The full operon is referred to as the itaconate response operon (IRO) and is under the control of the LysR-family transcription factor, ItaR [35,36]. This is the first operon of *Salmonella* pathogenicity island (SPI)-13 [37,38], and the IRO is one of the most upregulated operons during STm invasion of macrophages [39–41]. Additionally, various studies have shown that deletion of IRO genes in STm, in the related Enteritidis and Gallinarum serovars, or in *Yersinia pestis,* resulted in attenuated infection of macrophages [32,37,40–43] and mice [32,44–48]. While the importance of the IRO for infection has been demonstrated, we lack a clear understanding of how itaconate is used by STm during infection and the extent to which itaconate affects STm metabolism among the other stressors found within the SCV.

**Fig 1:**
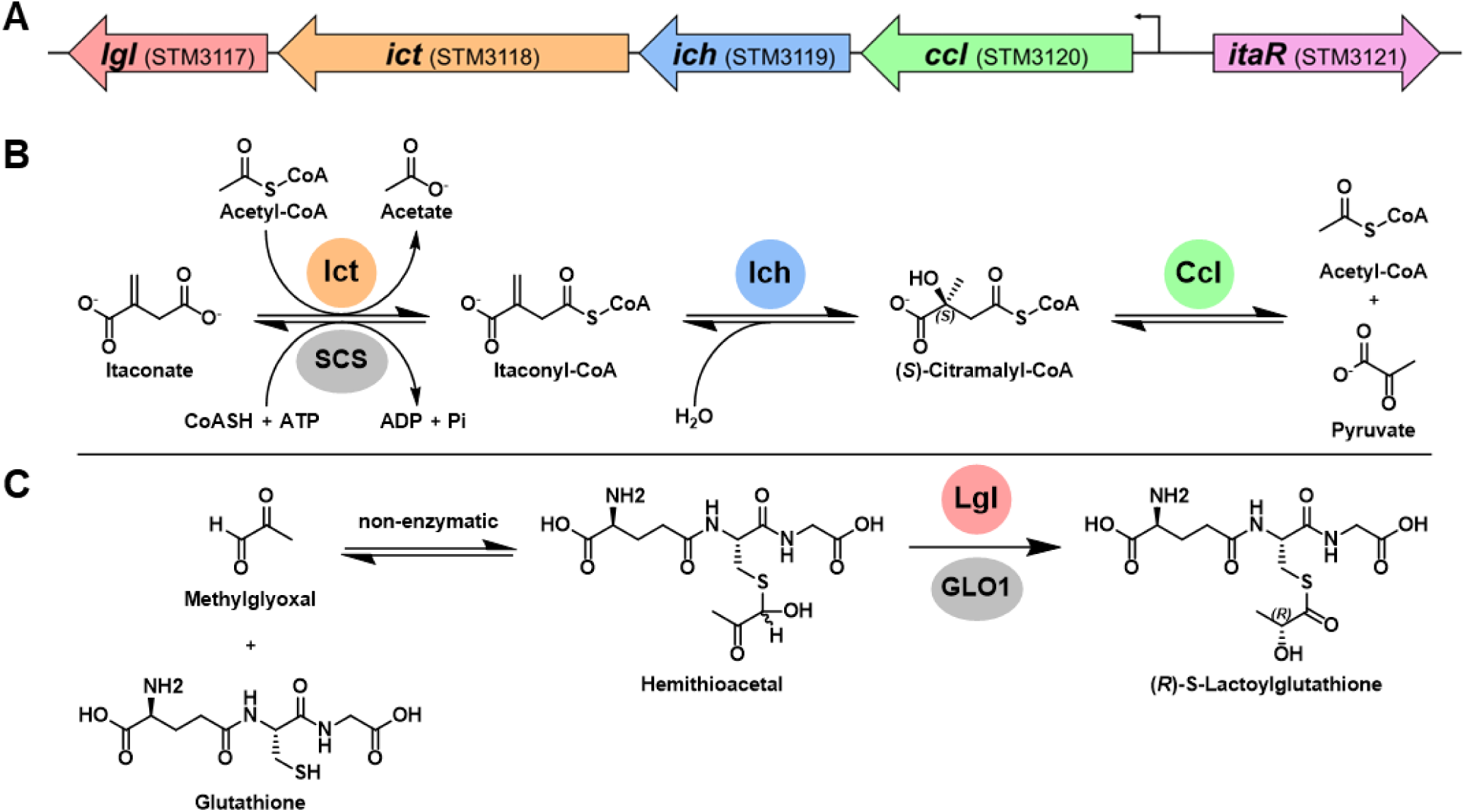
***S*. Typhimurium itaconate and methylglyoxal detoxification by the enzymes of the itaconate response operon (IRO).** (A) STm itaconate degradation enzymes (Ict, Ich, Ccl) are encoded on the SPI-13 together with lactoylglutathione lyase (Lgl), under the control of the itaconate regulator protein (ItaR). (B) Itaconate is degraded by three enzymes: Ict, Ich and Ccl. The action of Ict, a CoA-transferase, can be replaced by that of succinyl-CoA synthetase (SCS). (C) Lgl detoxifies the toxic metabolism byproduct methylglyoxal through isomerization of the spontaneously formed hemithioacetal intermediate. This reaction can also be performed by glyoxalase I (GLO1).

To add to our understanding of the effects of itaconate on STm overall, we decided to study STm growth on itaconate in axenic cultures using defined media. Expression of the IRO is not seen at any stage of growth in LB, a nutrient-rich media, nor in response to infection-relevant stress factors (*e.g.,* anaerobic, peroxide, nitric acid, bile shock), nor in other commonly used media that mimic the intra-macrophage environment [49]. In fact, the IRO has only been shown to be expressed in response to itaconate through ItaR activation [35,50,51]. As such, we chose to grow STm in the commonly used M9-minimial media with itaconate as the sole carbon source, referred to as M9I herein. We compared this with growth in the carbon-rich Mueller-Hinton broth (MHB), and another carbon-limited media, M9 with acetate as the sole carbon source (M9A). We found that STm growth in M9I is substantially slower than in M9A or MHB, but that the rate can increase with subsequent subcultures to adapt to itaconate. We identified two mutations that assist in this adaptation and used a metabolomics approach to map out the dominant effects on STm metabolism. We also found that Ich, Ccl, and ItaR are necessary for itaconate catabolism, but Ict is dispensable. In addition to improving our understanding of the interplay between STm and itaconate, our results suggest that the use of M9I can mimic the intracellular environment and could find application in the discovery of new antimicrobial molecules targeting nutrient stress [14].

## Results

### 1. STm growth on itaconate as a sole carbon source requires the ich, ccl, and itaR genes

To develop conditions in which we could study the itaconate degradation pathway and in turn, the effects of itaconate on STm, the bacterium was grown in M9I. STm can indeed grow in M9I, but at a drastically reduced growth rate compared to rich media (**Fig 2A**) [52,53]. Subsequent subcultures showed a consistent increase in the growth rate and OD at stationary phase. Comparing between the 2^nd^ and 6^th^ subcultures, the doubling time decreased from 11 h to 3 h, yet it did not reach that observed in MHB (0.4 h). The lag time decreased from ∼90 h to ∼40 h, compared to 1.8 h in MHB. Lastly, the maximum OD_600_ at stationary phase shifted from 0.54 in the 2^nd^ subculture to 1.7 in the 6^th^, compared to 2.5 in MHB.

**Fig 2:**
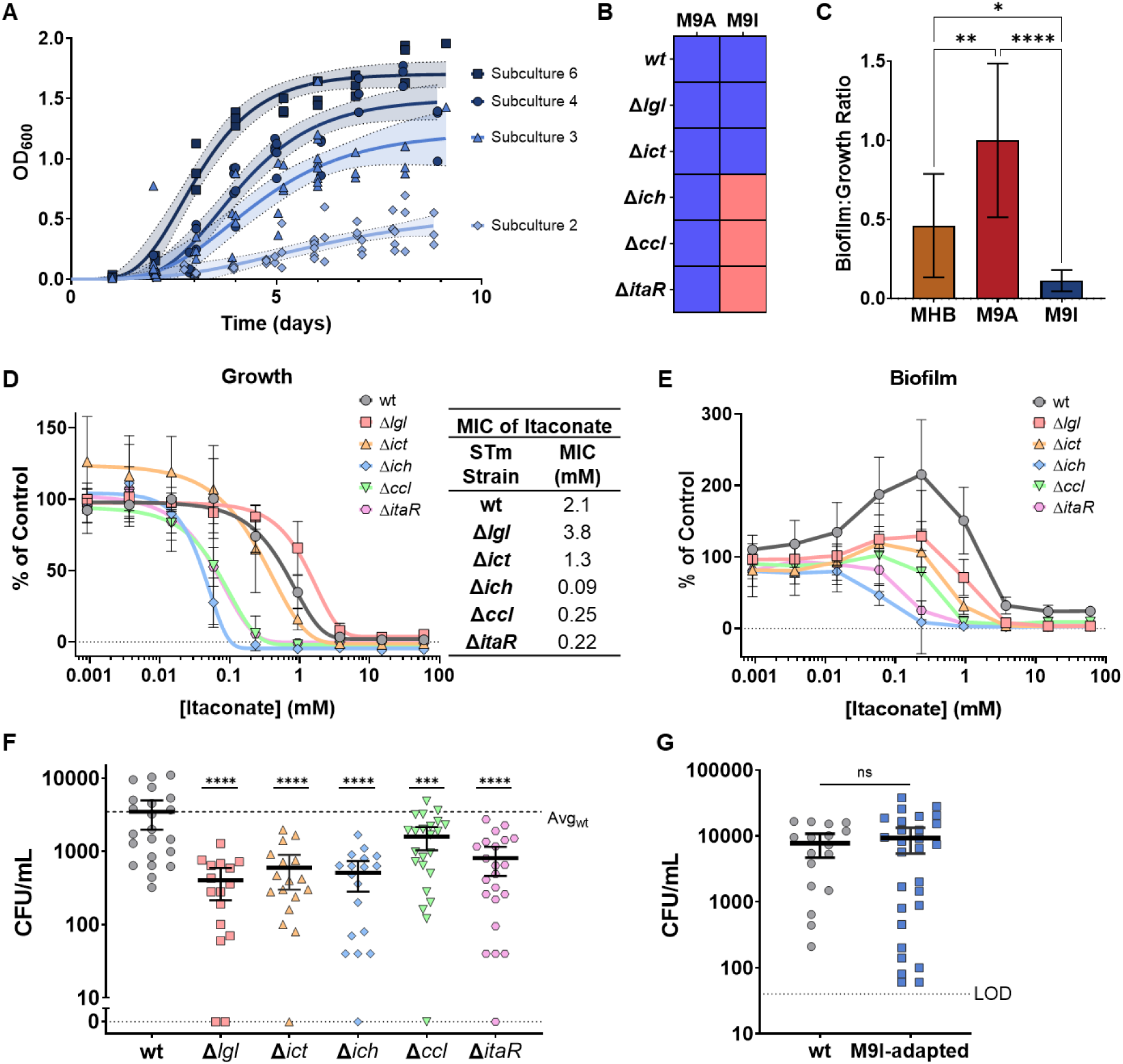
STm can grow in media using itaconate as a sole carbon source. (A) STm growth in M9I is initially slow, but after subsequent subcultures, it adapts and increases its growth rate. Logarithmic growth curves are shown with the bands denoting the standard error of each fit. (B) STm wt and IRO mutants grown in M9A and M9I, blue: growth and red: no growth. (C) Biofilm production by STm when grown in MHB, M9A, and M9I in 96-well plates. Biofilm was measured by crystal violet staining of the biomass, solubilizing and measuring the OD550. The measurements were normalized based on the planktonic growth (OD600) in each well. The data is given as a ratio of biofilm to planktonic growth; n ≥ 8, one-way ANOVA (with Tukey’s multiple comparisons test between groups). (D) Itaconate MICs for each of the STm IRO mutants measured in M9A. Data is presented as the percentage of growth (OD600) compared to the untreated control for each mutant, fit to a Gompertz MIC curve. Error bars denote the SD; n = 4. The MICs are tabulated. (E) Biofilm levels produced in the same itaconate MIC experiment as shown in D. The data is presented as a percentage of the amount of biofilm produced by the untreated control, for each mutant. Straight lines are used to connect the datapoints. (F and G) Intracellular survival of various STm mutants in RAW 264.7 macrophages, measured using a gentamicin protection assay. STm wt was compared with (F) the IRO mutants or (G) M9I-adapted STm. Error bars denote the 95 % CI of the mean. One-way ANOVA (with Dunnett’s multiple comparisons test with the wt) and two-tailed t-test used for comparing groups, respectively. All experiments were repeated a minimum of 3 times. *, p < 0.05; **, p < 0.01; ***, p < 0.001; ****, p < 0.0001; ns, non-significant, p > 0.05.

To understand which genes in STm are required for the catabolism of itaconate, single-gene deletion mutants of each of the IRO proteins were grown in M9I and compared with M9A as a negative control (**Figs 2B and S1**). Growth was observed for all mutants in M9A, while only wt, Δ*lgl* and Δ*ict* could grow in M9I. The cultures were left for 10 days without any visible growth for the Δ*ich*, Δ*ccl*, and Δ*itaR* mutants. This establishes that the Ich and Ccl enzymes, as well as the transcription factor, ItaR, are necessary for STm to degrade itaconate.

### 2. Biofilm production increases on acetate, but decreases on itaconate

Bacteria such as *P. aeruginosa* [54] and *S. aureus* [55] have been shown to respond to itaconate stress by increasing their production of biofilm. Thus, the amount of biofilm produced by STm when grown in MHB, M9A, and M9I was measured by crystal violet staining (**Fig 2C**). STm cultures were pre-adapted to the M9A or M9I media before measuring biofilm formation as described in the methods section. Interestingly, compared to the MHB culture, biofilm production was substantially increased in M9A, but decreased in M9I. An explanation for this is proposed later.

### 3. An inactive itaconate degradation pathway increases susceptibility to itaconate

STm has numerous mechanisms to respond to dicarboxylic acid stress, with the itaconate degradation pathway being the most direct one. To assess the effect that itaconate degradation has on the ability of STm to tolerate itaconate, we measured the MIC of itaconate for STm grown in M9A – a media in which the glyoxylate pathway and Icl (one of the enzyme targets of itaconate) are necessary. Consistent with the growth results presented in **Fig 2B**, the Δ*ich*, Δ*ccl*, and Δ*itaR* mutants (MIC of 0.09, 0.25, and 0.22 mM, respectively) were approximately an order of magnitude more susceptible to itaconate than wt STm or mutants Δ*lgl* and Δ*ict* (2.1, 3.8, and 1.3 mM, respectively; **Fig 2D**). Next, biofilm production was measured in response to itaconate treatment in M9A medium to see if STm and the IRO mutants exhibit any stimulation of biofilm production when itaconate is used as an antibiotic, rather than a carbon source. Indeed, wt STm was found to exhibit a large increase in biofilm at sub-MIC concentrations of itaconate, reaching a peak at ∼230 μM (**Fig 2E**). The biofilm response was dampened, though still observed, for Δ*lgl,* Δ*ict*, and Δ*ccl,* with the peak amount of biofilm being produced at ∼60 μM itaconate. Both the Δ*ich* and Δ*itaR* mutants did not appear to exhibit an increase in biofilm production at any concentration of itaconate. This suggests that having an intact itaconate degradation pathway—the resistance mechanism for the antibiotic being used here—is necessary to induce biofilm stimulation in response to itaconate.

### 4. STm IRO mutants are attenuated in their ability to infect macrophages

STm IRO mutants were previously shown to be attenuated in their ability to infect macrophages in screens of deletion mutant libraries [37,40–43]. These results were successfully reproduced by comparing each of the STm mutants in a gentamicin protection assay (in which the extracellular bacteria are selectively killed by gentamicin) using RAW 264.7 murine macrophages (**Fig 2F**). When compared to wt STm, all five mutants showed significantly attenuated intra-macrophage proliferation after an infection period of 20 h. Interestingly, the trends observed in the ability of different mutants to metabolize/resist itaconate did not translate to this experiment (see **Fig 2D**). Whereas wt STm had an average infection of 3500 CFU/mL, Δ*lgl* (400 CFU/mL) was the most attenuated, followed by Δ*ich* (510 CFU/mL), Δ*ict* (600 CFU/mL), Δ*itaR* (810 CFU/mL), and Δ*ccl* (1600 CFU/mL). Furthermore, the attenuation of Δ*ich* was observed 1 h after infection, after which the CFU was maintained over 72 h (**S2 Fig**). This seems to indicate that the diminished infection relates to an inability to colonize the macrophages.

### 5. Mutations in rpoS and dctA benefit growth on itaconate

To explore the ways in which STm adapts to more efficiently catabolize itaconate and improve its growth in M9I over subsequent subcultures, we sequenced the M9I-adapted and M9A-adapted STm strains. The same STm parent strain (cultured once in nutrient broth, NB) was subcultured three times in parallel in M9I and M9A. The dominant background mutations that were observed in the culture prior to M9I/A-adaptation (*i.e.* background differences between this parent strain and the ATCC 14028 reference genome) are listed in **Table S1**, and were filtered out before the following analysis of mutations identified in M9A and M9I. Across three replicates in each medium, 44 distinct mutations were observed for growth in M9A, with only six detected in all replicates (**S3A Fig**). Notably, none of the mutations found in the M9A-adapted strains were observed at a frequency above 25% and hence may not have conferred significant growth advantages. In the M9I cultures, 81 distinct mutations were noted, with an overlap of five genes across all replicates (S3B Fig). Further inspection of these mutations revealed that three of those five genes were also found to be mutated in the parent strain, but at different residues, and were also not found at frequencies above 25%. As such, those mutations could not be confidently linked to improved growth in M9I. The mutations identified in the remaining two genes were uniquely found in the M9I-adapted strains, in all three replicates, and were mutated in frequencies above 25% in at least one replicate. Thus, these likely conferred an advantage to growth in M9I (**Table 1**). The most abundant set of mutations were single nucleotide polymorphisms (SNPs) that introduced a stop codon into the gene of the alternative RNA polymerase sigma factor, *rpoS*. RpoS regulates a large group of genes during entry into stationary phase and as part of the general stress response of STm and other γ-proteobacteria, including a response to temperature, pH, osmolarity and oxidative stress [56]. Furthermore, RpoS has been established as an important virulence factor [57,58]. Four distinct nonsense mutations were observed across the three growth replicates at varying frequencies. If we presume that each bacterium contains only one of these SNPs, 95-100 % of each culture no longer had a functional RpoS. This strongly suggests that RpoS is a hindrance to growth in M9I.

**Table 1:**
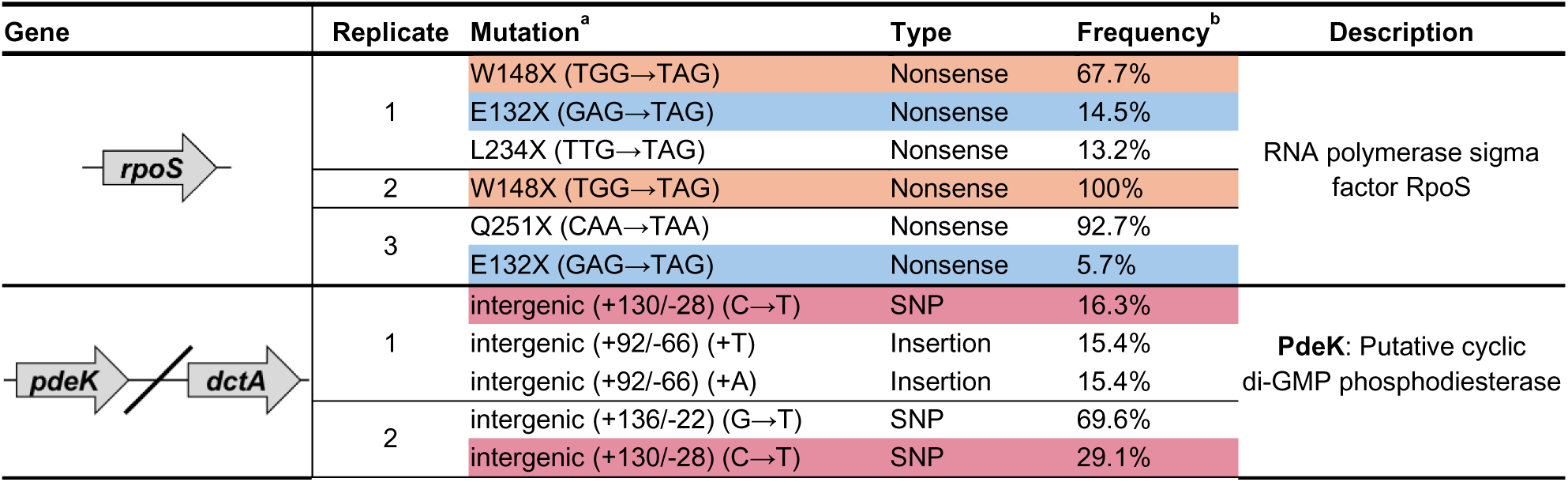
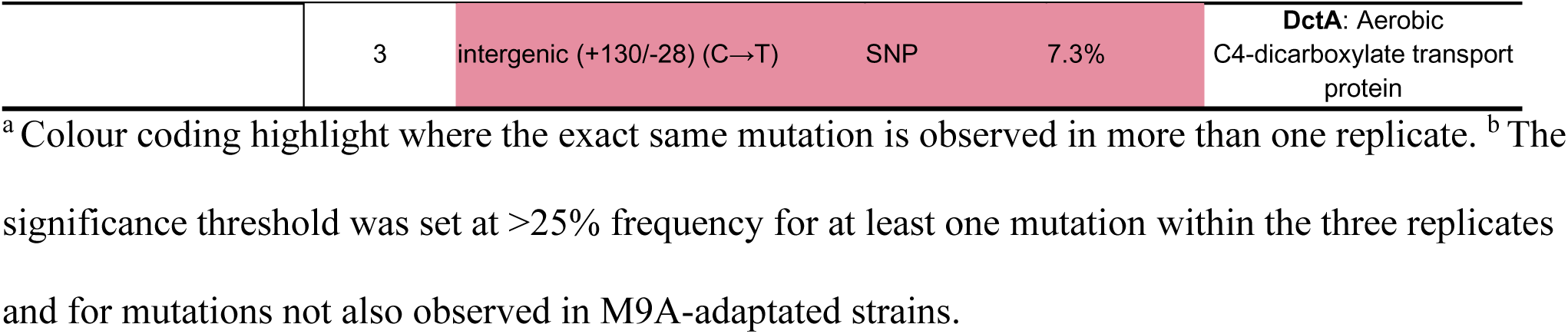
Significant STm genomic mutations during adaptation to M9I.

The second most abundant set of mutations were found in the intergenic region between the *pdeK* and *dctA* genes, upstream of *dctA* (**Fig 3**). PdeK is a putative cyclic di-GMP phosphodiesterase and DctA is an aerobic C4-dicarboxylate transport protein involved in importing molecules such as succinate, fumarate and malate into the cell [59]. Four mutations were identified in total, consisting of two distinct insertions (either A or T) directly before the -10 element of the promoter, and two SNPs in the 5’-UTR region of *dctA*, at an IscR transcription factor binding site [60]. One of the SNPs (C→T, +130/-28) was found in all three replicates, between 7-30 % frequency, while the other SNP (G→T, +136/-22) was found in only one replicate, although at the highest frequency (70%) among all replicates for this locus. Both insertion mutations were found in the same replicate at 15 % frequency. Based on their frequencies, the mutations in the *pdeK*/*dctA* intergenic region have a lesser impact compared to those of the *rpoS* SNPs, however, it is possible that over a few extra subcultures, higher frequencies in the *pdeK*/*dctA* mutations would arise. Taken together, these mutations in the *dctA* promoter region likely affect transcription levels. Based on previous studies identifying similar mutations [60,61] that are discussed later, we suggest that these result in higher expression levels of *dctA*, and increased transport of itaconate inside the cell.

**Fig 3:**
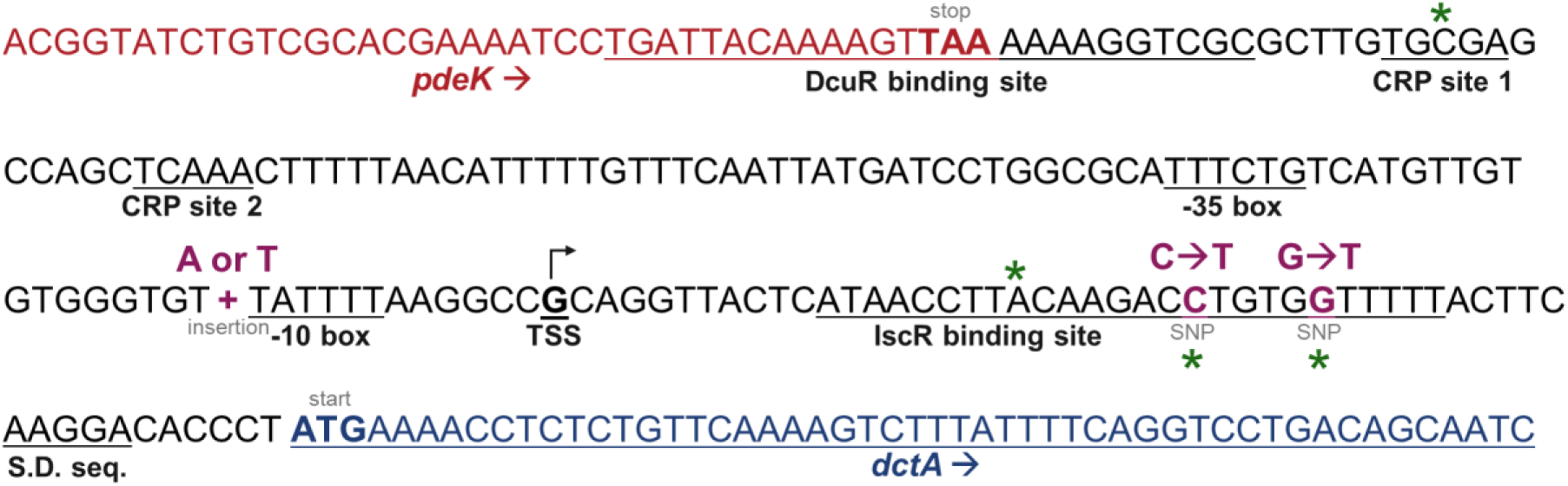
Mutations acquired selectively during growth in M9I that are found in the intergenic regions between *pdeK* and *dctA.* The two SNPs and two insertion mutations that were observed and descripted in Table 1 are in purple. Relevant transcription factor binding sites and promoter regions are annotated. Nucleotides denoted by a green * refer to sites for which mutations were previously characterized by Amin *et al.* (2022) [62] in the CRP binding region, or Quandt *et al.* (2014) [61] and Wenner *et al.* (2024) [60] in the IscR binding region, and that resulted in increased *dctA* expression and improved growth on C_4_-dicarboxylates. CRP: cAMP receptor protein, TSS: transcription start site, S.D.: Shine-Dalgarno sequence.

Given that RpoS is an important regulator of virulence for STm, the ability of M9I-adapted STm to infect macrophages was investigated. Previously, Δ*rpoS* mutants have been shown to have little to no attenuation in their ability to infect macrophages, but were attenuated in mice models [63]. To confirm this phenotype, and determine if the other mutations factored in, proliferation of the wt and M9I-adapted STm were compared in RAW 264.7 macrophages (**Fig 2G**). Indeed, no significant difference between the means of the two strains was observed. The wt STm burden was 7700 ± 3100 (95% CI) CFU/mL and the burden of the M9I-adapted strain was 9300 ± 3900 (95% CI) CFU/mL. Altogether, this suggests that the mutations acquired to improve itaconate catabolism do not affect the ability of STm to infect macrophages.

### 6. Remodeling of STm metabolism when grown on itaconate

To gain further insight into the general adaptations of STm to itaconate, a LC-MS-based targeted metabolomic analysis was performed. The metabolism of STm was compared at both a mid-exponential phase (EP) and at an early-stationary phase (SP) of growth in MHB, M9A, and M9I. STm growth curves were first reestablished under the larger-scale conditions used for the metabolomics experiment to appropriately determine the OD at the EP and SP endpoints. (**Fig 4A**). The detection methods used were optimized to search for a panel of 296 ions in total, and overall, 129 ions were quantified and included in the analysis. Principal component analysis (PCA) revealed that the metabolic profiles of STm grown in each medium differed considerably, while the profiles at different stages of growth (EP versus SP) within each medium were more similar (**Fig 4B**). The M9I EP and SP profiles completely overlapped, while only one M9A SP sample fell outside of the EP 95 % confidence region. The largest difference between growth phase profiles was seen by STm grown in MHB, although the difference was small compared to the profile differences between media. The significance of metabolite differences across all conditions were determined by two-way ANOVA, summarized here as a heatmap showing the relative abundances of the 40 most significant (ranked by p-value) metabolites (**Fig 4C**). An examination of the pathway ontology revealed that the nucleotide and carbohydrate metabolic pathways were the most abundant among significant features (**Fig 4D**). A secondary analysis consisting of pairwise comparisons (T-tests) between media conditions (stratified by growth phase) was also done to identify the specific fold changes (FC) in significant (log_10_(FC) > 2 and p-value < 0.05) metabolites across the different condition sets (**S4A Fig**). Multiple pathway enrichment analysis using both the ANOVA and T-test/FC identified significant metabolites and were used to guide our understanding of these results (**S4B-D Fig**). However, no specific pathways were found to be markedly (p < 0.05) enriched by this method, which appears to indicate that the effects across the three media are on the STm metabolism as a whole. As such, the pathways most abundant in significant metabolites, as determined by two-way ANOVA across all six conditions, were prioritized for analysis since they could give a clearer picture of metabolic differences across the three media.

**Fig 4:**
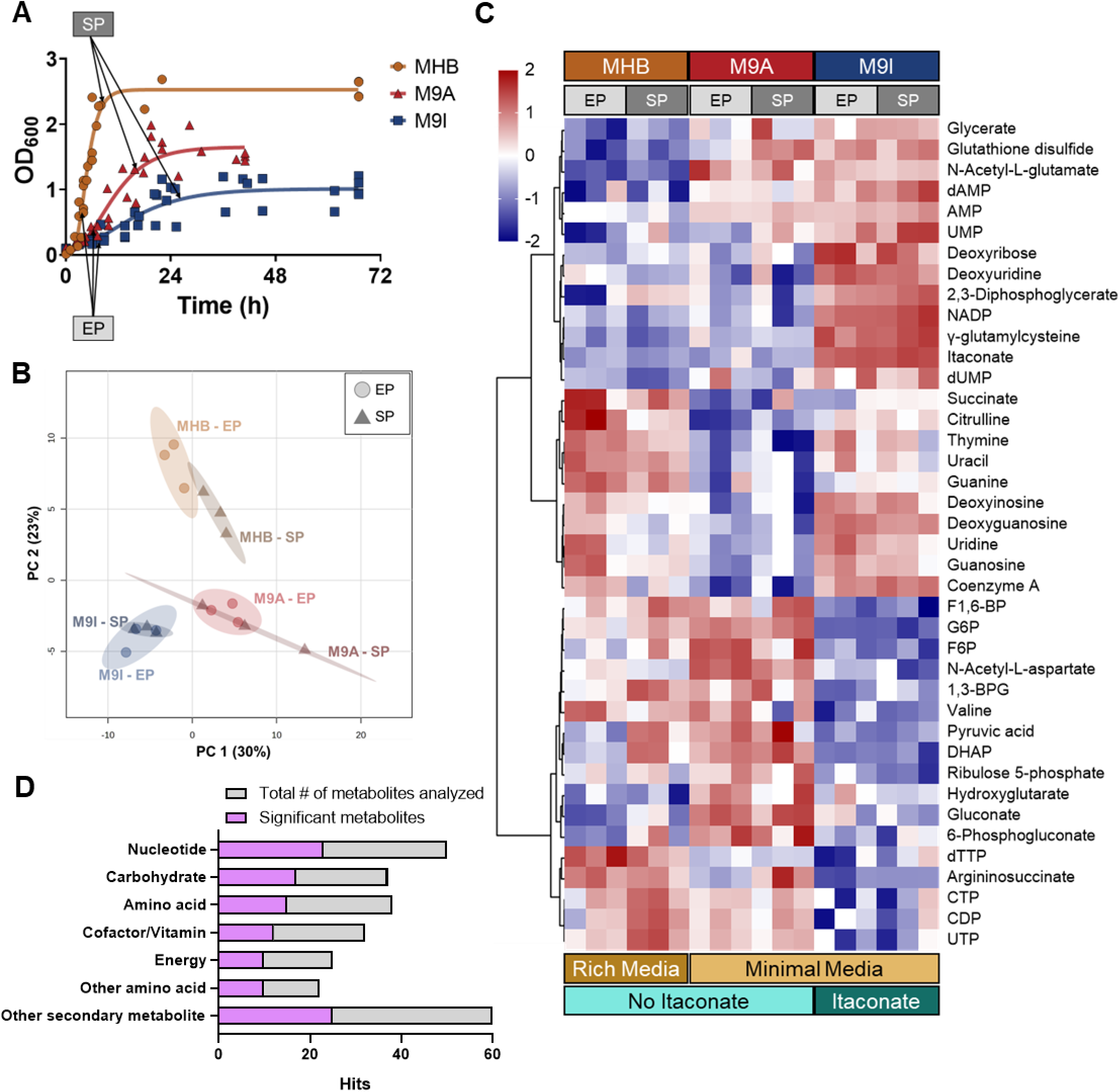
Metabolomics of STm in MHB, M9A, and M9I at two phases of growth. (A) Growth curves in each medium under the culture conditions used for metabolomics with the mid-exponential phase (EP) and early-stationary phase (SP) identified. (B) Principle-component analysis of all six conditions. Ellipses indicate 95% CI, n = 3 per condition. (C) Heat map showing the normalized abundance of each of the most significant metabolites as identified by a two-way ANOVA (ranked by p-value and using Ward’s method to cluster features). (D) Pathway ontology analysis showing both the total number of metabolites measured by our HPLC-MS method and those that are significant in each pathway.

***i) Effects on carbon metabolism.*** When growing on non-glycolytic carbon sources such as acetate or succinate, STm and other enteric bacteria rely on gluconeogenesis for the anabolic production of the various sugars necessary for growth and cellular functions, such as cell wall and nucleotide biosynthesis [64–66]. In M9A, the metabolomics analysis reveals an abundance of gluconeogenic metabolites, as well as pentose phosphate pathway (PPP) sugars, consistent with active growth (**Fig 5**). However, when grown on itaconate, the opposite trend is observed, with a broad and drastic decrease in metabolites within the glycolysis/gluconeogenesis and PPP pathways. Given that itaconate enters carbon metabolism in the form of acetyl-CoA and pyruvate after being degraded, a similar need for gluconeogenesis would be expected when growing in M9I. The reduced flux in this pathway could be a factor in the substantially reduced growth rates of STm in M9I.

**Fig 5:**
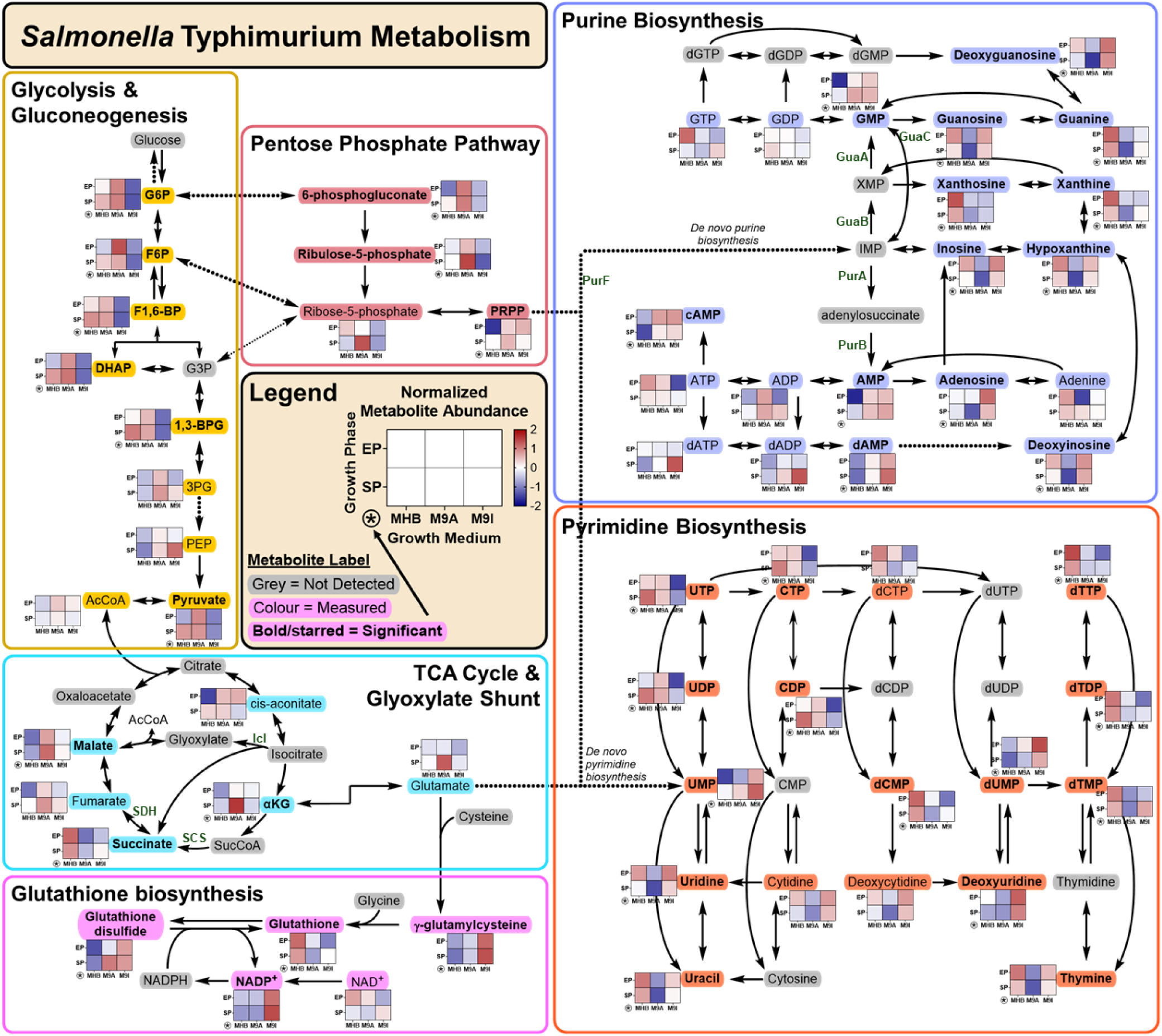
Pathway analysis of STm metabolomic results. Relative metabolite abundances across the three media (MHB, M9A, M9I) and two growth phases (EP and SP) are shown as a single heatmap for selected metabolites quantified in our method and placed adjacent to the molecule in the pathways. Double sided arrows indicate reversible reactions performed by the same enzyme, double arrows in opposite directions indicate reversible processes that are performed by different enzymes. Dotted lines indicate multiple enzymatic steps. G6P: glucose-6-phosphate, F6P: fructose-6-phosphate, F1,6-BP: fructose-1,6-bisphosphate, DHAP: dihydroxyacetone-phosphate, G3P: glyceraldehyde-3-phosphate, 1,3-BPG: 1,3-bisphosphoglycerate, 3PG: 3-phosphoglycerate, PEP: phosphoenolpyruvate, AcCoA: acetyl-CoA, PRPP: 5-phosphoribosyl-1-pyrophosphate, αKG: α-ketoglutarate, SucCoA: succinyl-CoA.

For the TCA cycle, our results are less clear because no significant accumulation or depletion of any detected TCA metabolite is seen for STm in M9I. Itaconate is a known inhibitor of at least three enzymes in the STm TCA and glyoxylate cycles: Icl, SDH, and SCS. Inhibition of Icl cannot be confidently established because isocitrate could not be measured. However, the slight increase in abundance of cis-aconitate during the M9I exponential phase is consistent with Icl inhibition. This is similarly the case for SCS, which converts SucCoA to succinate; SucCoA was not measured, and little information can be gleaned from the levels of α-ketoglutarate (one metabolite before SCS) in M9I. Furthermore, the lack of succinate accumulation suggests that any SDH inhibition is not affecting the flux through this part of the cycle compared to the two other media. In M9A, the observed accumulation of malate could indicate activity through the glyoxylate cycle, which is important for growth on acetate [67].

***ii) Effects on glutathione metabolism.*** Exposure to electrophilic compounds, such as itaconate, is known to induce specialized electrophilic stress responses [68]. Glutathione (GSH) is an antioxidant that is often employed in response to electrophilic and oxidative stress. In the case of itaconate, it has been shown to form adducts through thia-Michael addition to itaconate [69]. By comparing GSH levels to those of its oxidized pair, glutathione disulfide (GSSG), we can infer the oxidative state of the cells under each condition (**Fig 5**). When growing on itaconate, the GSH to GSSG ratio strongly favours the oxidized form. This is true also for growth in M9A at stationary phase, but the ratio is flipped in MHB, indicating little or no oxidative stress in this rich media. In M9I, we also see an abundance of the glutathione precursor γ-glutamylcysteine, which may indicate a general upregulation of glutathione production in response to this stress. Similarly, NADP^+^, the oxidized form in the NADP^+^/NADPH redox pair required by enzymes for cycling between GSH and GSSG, is also found in relatively large quantities in M9I compared to both MHB and M9A. Although it is difficult to draw conclusions from the NADP^+^ abundance alone, the high amounts detected support the observation of elevated oxidative stress in M9I, with NADPH recycling being slower than GSSG formation.

***iii) Effects on nucleotide biosynthesis.*** Given that nucleotide biosynthesis was the most represented pathway in our data and that recently, itaconate was found to affect the *de novo* biosynthesis of purines in STm [24] and Mtb [25], the effects of itaconate on purine and pyrimidine biosynthesis were explored (**Figure 5**). To our discontent, the molecules involved in *de novo* purine biosynthesis were not detected, thus limiting our ability to compare with previous results. Nonetheless, trends in the broader metabolic profiles for the pathways downstream of the *de novo* portion of purine and pyrimidine biosynthesis were identified. The previous work had observed decreases in the end products of *de novo* purine biosynthesis (IMP, AMP, and GMP) in response to itaconate treatment through inhibition of the first step in the pathway, PurF, and the GuaABC enzymes in GMP biosynthesis [24,25]. The data presented here shows that the GMP and AMP levels are not depleted for cultures in M9I, and are at similar levels in M9A, whereas the levels were considerably lower in MHB. Considering that we are analyzing the metabolomes of media-adapted STm that have established a new homeostasis and growth rate within each media (compared to experiments in which itaconate exposure is a treatment group), it follows that we are likely not to observe the same responses as in prior studies.

Broadly, in M9I, we observe a depletion in the supply of both purine and pyrimidine nucleotide di-and triphosphates that is particularly prominent in the exponential phase compared to stationary. This could indicate a general lack in energy supply, or a lag in the ability of STm to replenish nucleotide triphosphate stores needed for transcription at a pace that matches the growth rate in this phase. In the case of deoxynucleotides, we see an accumulation of all three species of adenine phosphate (dAMP, dADP and dATP) in the stationary phase. For the pyrimidine deoxynucleotides, we generally see lower quantities of these species in M9I, however the trend of abundance increasing from EP to SP remains. This accumulation in SP could be the result of a reduced ability to downregulate the synthesis of deoxynucleotides in M9I once the cells stop actively growing. Furthermore, we see increased levels of purine and pyrimidine deoxynucleosides and some nucleosides in M9I, especially when compared to M9A for which a general deficit of these species is observed. This could be related to usage of the salvage pathways. *De novo* purine and pyrimidine biosynthesis stems from the metabolite phosphoribosyl diphosphate (PRPP) in the PPP. Given the higher flux through PPP that we observed in M9A, STm may be better able to rely on *de novo* biosynthesis of nucleotides under these conditions. However, in M9I, given the lack of gluconeogenesis observed, the bacteria may rely more on the salvage pathways for their nucleotides.

Another metabolite of interest is cyclic-AMP (cAMP), which is a secondary signalling molecule that regulates transcription through the cAMP receptor protein (CRP) in response to changes in carbon source, especially in the absence of glucose [70]. As expected, a higher abundance of cAMP was seen in both M9I and M9A compared to MHB.

## Discussion

We have shown that STm can grow on itaconate as a sole carbon source, and that growth rates on itaconate increase with subculturing. Itaconate degradation requires Ich, Ccl, and the IRO regulator ItaR to accomplish this. The first enzyme in the itaconate degradation pathway, Ict, is dispensable, although the growth of STm Δ*ict* in M9I is still attenuated. Similarly, the other enzyme in the IRO, Lgl, was not required for itaconate degradation, as expected given its function in methylglyoxal detoxification. These findings correlate with itaconate susceptibility in M9A, in which a ∼10-fold decrease in MIC was observed for STm mutants lacking one of the three required IRO genes compared. This is the first report identifying the genes of the STm IRO that are necessary for itaconate degradation, and is consistent with prior work on itaconate degradation in *P. aeruginosa*. In *P. aeruginosa* Ich and Ccl are required for growth on itaconate [20,71–73], with some groups reporting that Ict deletion prevents growth up to 24 h [20,54] or 36 h [72], while another saw attenuated growth after 36 h in a *Pseudomonas putida* strain heterologously expressing the *P. aeruginosa* IRO [71]. It may be that *P. putida,* and not *P. aeruginosa,* has an Ict alternative (as seen with STm) that allows growth without Ict. However, longer growths of Δ*ict P. aeruginosa* on itaconate are needed to confirm this. To our knowledge, *P. aeruginosa itaR* deletions mutants have not been tested in their ability to degrade itaconate. Furthermore, we were able to corroborate previous work showing that each single gene deletion mutant of the STm IRO genes results in a reduced ability to infect macrophages, regardless of their necessity for degradation [37,40–43].

The extended lag phase observed in M9I and subsequent adaptation to the M9I media seem to indicate that itaconate is not a preferred carbon source for STm under these conditions. The nonsense mutations introduced into *rpoS* to near fixation after only three subcultures is indicative of a RpoS-induced inhibition of itaconate catabolism, uptake, or otherwise. RpoS is a generalized stress response and a stationary phase sigma factor that competes with the primary sigma factor RpoD and others for the core RNA-polymerase [56]. As such, high RpoS levels results in reduced growth through RpoD competition. When this is detrimental to bacteria, RpoS mutations are frequently selected for to correct this through antagonistic pleiotropy, which is often reversible [74,75]. Our finding that M9I adaptation did not affect STm infection of macrophages is consistent with the previous finding that *rpoS* deletion does not affect infection or survivability in macrophages *in vitro* [63]. Furthermore, RpoS has been linked to biofilm formation through multiple transcription regulators [76–78]. The observed increased baseline biofilm levels between cultures in M9A compared with M9I—along with the fact that wt STm grown in M9A is stimulated to produce excess biofilm by sub-MIC concentrations of itaconate—leads us to believe that the *rpoS* mutations are a factor in the reduced biofilm production phenotype observed in M9I growth. Biofilm production when grown on non-glycolytic carbon sources is supported by gluconeogenesis [79]. As such, we can correlate the higher biofilm levels in M9A and the lower levels in M9I, with their respective increase and decrease in metabolic flux through gluconeogenesis and PPP observed in our metabolomics results.

A reduction in biofilm stimulation at sub-MICs of itaconate was also detected for the IRO mutants, particularly those incapable of degrading itaconate. The sub-MIC biofilm stimulatory response is common among biofilm-forming bacteria as part of their stress response to antibiotic treatment, with some differences observed between classes of antibiotics [80]. Recently, in *E. coli,* this phenomena has been directly linked with global changes in central metabolism and respiration in response to antibiotic treatment [81]. Yaeger *et al.* [81] showed that knockout mutants of the TCA cycle (among others) have a hampered biofilm stimulation response to sub-MIC antibiotic stress. Given that itaconate directly inhibits multiple enzymes in the TCA cycle, it follows that limiting/eliminating the ability of STm to detoxify itaconate in the knockouts results in more TCA cycle inhibition and a similar decrease in biofilm production as seen with the *E. coli* TCA-cycle mutants. The observation of STm Δ*lgl* reduced biofilm formation can be included in this hypothesis through a reduced ability to handle itaconate-induced oxidative stress via reduced methylglyoxal detoxification and its downstream effects [33]. Complicating this hypothesis is a study that looked at itaconate treatment of STm in LB cultures, in which they did not observe any sub-MIC increase in biofilm production [82]. This will require further investigation to properly understand the STm biofilm response to itaconate treatment, how growth on different carbon sources affects it, and how it relates to the biofilm stimulation by itaconate seen in other species [54,55].

Various C_4_-dicarboxylates (C4DCs; *e.g.* succinate, fumarate, L-aspartate) are taken up and used by STm and other enteric pathogens as carbon sources or signaling molecules at various stages of infection [83,84]. A key example is succinate, a structural analog of itaconate, which is used by STm to fuel its growth during infection of the gut, where succinate is produced by the host microbiota [85]. However, during infection of macrophages, STm uses glucose and other glycolytic intermediates as a preferred carbon source over succinate [60,86–88]. This is despite the fact that macrophages accumulate succinate during infection due to a shift in the host cell metabolism from oxidative phosphorylation to glycolysis [86,89]. Instead, this succinate is recognized by *Salmonella* as a molecular signal to induce virulence genes important for intracellular survival through expression of SPI-2 [90]. Succinate and other C4DCs are taken up aerobically by the DctA transporter and anaerobically by DcuB [84,91]. DctA has also been shown to be sufficient for itaconate transport [92]. RpoS has been reported to restrict STm uptake and catabolism of C4DCs through indirect repression of *dctA* [60,93], and direct repression of TCA cycle genes used for C4DC catabolism [94,95]. STm growth on succinate exhibits an extended lag phase, even compared to other enteric bacteria like *E. coli*, and deletion of *rpoS* was also shown to substantially shorten the lag phase [93]. In light of this, we can reasonably conclude that *rpoS* suppression of itaconate uptake and metabolism genes is the reason for the rapid selection of non-functional *rpoS* mutants during growth in M9I.

The mutations observed in the *dctA* promoter region are also consistent with our understanding of *dctA* regulation through both RpoS and non-RpoS mechanisms. In fact, the transcriptional regulation of *dctA* is known to be controlled directly by at least three transcription factors: the membrane-bound DcuS-DcuR two-component system, the cAMP receptor protein (CRP), and the iron sensing regulatory protein, IscR (**Fig 3**) [60,83,91]. CRP senses cAMP increases under carbon-limiting conditions, leading to induction of many genes, including *dctA* [70]. It is also responsible for catabolite repression of *dctA* expression when preferred (*i.e.* glycolytic) carbon sources are available. Recently, a SNP in the CRP-binding site of the STm *dctA* promoter region was reported to increase CRP induction of DctA, allowing growth on normally non-permissibly low orotate concentrations (which is also transported by DctA) [62]. On the other hand, IscR represses *dctA* expression and inhibits growth on succinate [60]. Previously, in two separate studies, a total of three SNPs in the IscR binding site of the *dctA* 5’-UTR resulted in reduced IscR-mediated repression, increased *dctA* expression, and an improved ability of both STm and *E. coli* to grow on succinate or other C4DCs [60,61]. To our satisfaction, both of the SNPs identified here through M9I-adaptation coincided with those previously discovered. The C→T (+130/-28) mutation is the exact same as that found by Wenner *et al.*,[60] and the G→T (+136/-22) SNP involves the same G nucleotide reported by Quandt *et al.* [61], although they observed a transition to an A, and we observed a transversion to a T. In reference to the A or T insertion observed immediately upstream of the *dctA* -10 element in our study, it has been shown that STm RpoD has a higher tolerance for an “extended” -10 element compared to *E. coli* [96]. These results suggest that this last mutation may result in a positive, if not neutral, effect on RpoD binding to promote *dctA* expression. In summary, both sets of mutations acquired during M9I-adaptation are expected to translate into an improved ability of STm to uptake itaconate through DctA and a reduced suppression of the TCA cycle genes necessary for growth on C4DCs.

Recalling that many proteins have been identified as targets of *S*-itaconation, it was interesting to not find any mutations in our genotyping results for cysteines to another amino acid. While we would not expect active site cysteine residues of enzymes to be mutated, any instances of negative allosteric effects due to *S*-itaconation would be a possible site for mutation to prevent that. However, even when looking at the low frequency mutations, there were only 2 instances of cysteines being mutated to another amino acid, while in 8 cases, cysteines were gained. This finding suggests that in STm, most *S*-itaconation sites that are not in an enzyme active site may have a neutral or positive effect on enzyme function, such that mutation of the cysteines was not beneficial to the cells during growth on itaconate. This is consistent with the fact that in prokaryotes, all the validated targets of *S*-itaconation inhibition have been within enzyme active sites [18], whereas in eukaryotic cells, there are many instances of *S*-itaconations at regulatory sites resulting in altered protein functions. Alternatively, it could be that STm employs inducible protective strategies to sufficiently protect reactive cysteines, whether through glutathione-mediated protection or other unexplored mechanisms. For example, *S. aureus* has been shown to employ reversible *S*-bacillithiolations of protein cysteines in response to itaconate-and succinate-induced stress. However, STm is not known to make bacillithiol.

In light of the mutational adaptations uncovered herein, it is crucial to consider the potentially far-reaching effects that these may have when interpreting metabolomic results. Without an RpoS stress response, STm must rely on non-RpoS mechanisms to manage the stress of prolonged exposure to itaconate during growth in M9I. A key indicator of this stress was the increased flux through the glutathione metabolism pathway. Higher levels of both γ-glutamylcysteine and GSSG, point towards a greater need for glutathione during growth on itaconate. The lower levels of GSH detected in M9I compared to the other media are likely due to the sequestration of GSH as oxidized derivatives. It is known that GSH can form adducts with itaconate in mammalian cells as a means of buffering electrophilic stress [69]. This has been seen with fumarate, as well, with the fumarate-GSH adducts being recycled by glutathione reductase, the enzyme that uses NADPH to reduce GSSG back into GSH [97,98]. However, no such recycling pathway for itaconate-GSH adducts has been identified. It is possible that there is a population of itaconate-GSH adducts that accumulates in bacteria, and more research will be needed to explore the presence of these adducts in STm and whether the cells are able to recycle the adducts. A similar phenomenon was seen with STm during exposure to peroxide in glucose-rich medium that resulted in excess production of methylglyoxal [99]. Activation of the methylglyoxal detoxification pathway created a glutathione sink, reducing the available GSH levels. Interestingly, this connects Lgl activity as a homolog of GLO1 with the rest of the IRO, both of which respond to different aspects of oxidative stress found in the SCV.

Previous studies have explored the effect of itaconate on the metabolism of various bacteria when grown in rich media including *E. coli* (grown in LB) [82], Mtb (7H9) [25], *P. aeruginosa* (LB) [20], *V. cholerae* (LB) [100], and *S. aureus* (RPMI or MHB) [101,102]. In each study, flux through the TCA cycle was highlighted, with observations generally consistent with the known targets of itaconate inhibition. These studies carry the context that they were evaluating growth disruption by itaconate in carbon-rich media. As such, our results are distinguished from these by exploring the overall impact on STm metabolism and for STm growing on itaconate, rather than the ways in which it affects the metabolism of other carbon sources.

Of note, *P. aeruginosa* was shown to be able to efficiently catabolize itaconate, incorporate it into its central metabolism and utilize it for biofilm production through gluconeogenesis [20]. This is consistent with the fact that *P. aeruginosa* has a different itaconate degradation pathway from STm [26] and more readily proliferates on itaconate as a sole carbon source [20,54,72]. In addition to the low homology between *P. aeruginosa* Ict/Ich enzymes and those of STm [26], *P. aeruginosa* also has a dedicated itaconate transporter found within its IRO. This transporter was reported to be necessary for efficient growth on itaconate [71–73]. This can be contrasted with our findings that STm required mutations to increase DctA transporter expression, and elimination of RpoS-induced repression of C4DC usage in central metabolism. Taken together, it appears that *P. aeruginosa* is better equipped for use of itaconate as a carbon source compared to STm and likely other species with homologous IROs.

The full extent to which *Salmonella* uses itaconate as a carbon source or molecular signal during infection is highly relevant to the development of antibiotics that target intracellular bacteria. Herein, more light is shed on this important phenomenon. STm exhibits an extended lag phase and extremely slow growth rate in M9I that is ameliorated by rendering it RpoS non-functional and increasing itaconate uptake via augmented DctA expression. This is paralleled by STm use of succinate as a carbon source, for which slow axenic growth is improved through deletion of *rpoS* or introduction of mutations that improve DctA expression [60,93]. This points towards itaconate likely playing a similar role to succinate in the macrophage [60], where it acts as an important signalling molecule rather than as a carbon source. This is supported by both the shared transport machinery and catabolite repression mechanisms of the two molecules. Despite this, the importance of an intact itaconate degradation pathway during infection is clear and continues to be shown in studies; suggesting that it is a valid target for the development of novel antibacterials, which was the goal of one previous study [103]. The effects of itaconate under other infection-relevant conditions, such as at low pH—where itaconate has been shown to have more potent antibacterial activity—is an area of study that may also reveal considerably different metabolic effects [17]. This work has expanded on our understanding of STm itaconate degradation and the important interplay with other stress responses. Furthermore, using a media with itaconate as a sole carbon source enables high throughput screening of potential itaconate degradation inhibitors as an alternative treatment against intra-macrophage STm or other itaconate degrading bacteria.

## Methods

### Chemicals, media and bacteria strains

All chemicals, BD Difco™ Nutrient broth/agar (NB/NA), and Mueller-Hinton broth II (MHB) were purchased from Millipore-Sigma (Canada), Thermo Fisher Scientific (Canada) or Chem-Impex (USA) unless otherwise stated. M9 media was prepared from the following components: M9 salts (48 mM Na_2_HPO_4_, 22 mM KH_2_PO_4_, 8.6 mM NaCl, 18.7 mM NH_4_Cl), 2 mM MgSO_4_, 0.1 mM CaCl_2_, trace metals (30 μM FeCl_3_, 25 μM Na_2_EDTA, 8 μM ZnCl_2_, 3 μM H_3_BO_3_, 3 μM MnCl_2_, 2 μM (NH_4_)_6_Mo_7_O_24_, 2 μM CuCl_2_, 2 μM CoCl_2_), and either 0.4% (w/v) sodium acetate for M9A, or 10 mM itaconic acid for M9I. The media was adjusted to pH 7.2 before filter sterilizing.

*Salmonella* Typhimurium ATCC 14028, *Pseudomonas aeruginosa* ATCC 27853, and RAW 264.7 (TIB-71) were purchased from Cedarlane (Canada), and STm mutants (Δ*lgl*, Δ*ict,* Δ*ich,* Δ*ccl,* Δ*itaR*) are from the BEI Resources (www.beiresources.org) collection of single gene deletion (SGD) mutants that were originally constructed by Porwollik *et al.* (2014) [104].

### General bacterial growth procedures

Bacteria were routinely grown by first streaking from a glycerol stock on nutrient agar plates (with 50 μg/mL kanamycin for the STm mutants) and grown overnight at 37⁰C. Generally, 3-5 colonies were picked and transferred to 5 mL liquid cultures using the appropriate broth and antibiotic. Cultures were grown overnight at 37⁰C and 250 rpm. Stationary phase cultures were subcultured at a minimum 1/100 dilution into the appropriate fresh media and grown to a mid-exponential phase (OD ≈ 0.3-0.7) before use in experiments. Bacteria grown in minimal media were subcultured additional times (2 subcultures minimum in M9A and 3 subcultures minimum in M9I, unless otherwise stated) before use in experiments to allow for the bacteria to sufficiently adapt to the media. ODs were monitored using either a Cary 60 UV-Vis spectrophotometer (Agilent Technologies) or a SpectraMax i3x microplate reader (Molecular Devices). Bacterial growth curves were fit to Gompertz growth models using Graphpad Prism v9.0 (Graphpad Software) and doubling times were extracted using the Dashing Growth Curves[105] web application: https://dashing-growth-curves.ethz.ch/.

### Minimum inhibitory concentration (MIC)

MIC assays were performed following the microplate dilution antimicrobial susceptibility testing outlined in the CLSI guidelines with a few modifications [106]. Briefly, itaconic acid was dissolved to 1 M in M9A media and the pH was adjusted back to pH 7.2 to ensure no effect from pH changes as previously reported [17]. Serial dilutions were made in 96-well plates at the desired concentrations. Mid-exponential phase M9A-adapted bacteria were added at 5 × 10^5^ CFU/mL and the plates were sealed with a gas permeable moisture barrier seal (FroggaBio, Canada) to prevent excessive evaporation, before incubation at 37⁰C and 350 rpm, using a tabletop orbital plate shaker (OHAUS, USA). Growth was measured at OD 600 by microplate reader. The plate was monitored until there was visible growth in the untreated control (OD_600_ > 0.2, ∼36 h). The data was fit to a Gompertz non-linear regression [107], allowing for calculation of the MIC using Graphpad Prism v9.0. All MICs were done in a biological triplicate.

### Biofilm assays

Biofilm formation was quantified in 96-well plates using the methods established by Coffey and Anderson (2014) [108]. After either growth or MIC measurement, the suspended bacteria were first removed from the plate, before washing the plate twice with excess H_2_O, and staining the plate with 200 µL of 0.1% (w/v) crystal violet for 10 min at room temperature (RT). The plates were then rinsed four times with excess H_2_O, before allowing them to dry at RT overnight. Once dry, the stained biofilm was solubilized in 200 µL 30% acetic acid, incubated with shaking (350 rpm) for 10 min at RT, and the absorbance was measured at 550 nm. Biofilm formation in MHB, M9A, and M9I was measured at stationary phase (OD_600_ ∼0.35, ∼0.32, and ∼0.19, respectively in 100 μL/well), and the experiment was done in triplicate.

### Macrophage infection assay

Raw 264.7 mouse macrophages were cultured in Dulbecco’s Modified Eagle Medium (DMEM) with 10% fetal bovine serum (FBS) and grown at 37⁰C in a 5% CO_2_ atmosphere. A cell scraper was used to lift cells off the flask when passaging (rather than trypsin). Macrophages were seeded in 96-well plates at 1 x 10^4^ cells per well (100 μL; measured by haemocytometer) and allowed to adhere for ∼16 h before the experiment. On the day of the experiment, STm strains were subcultured 1/100 in NB and allowed to grow to mid-exponential phase. The culture was diluted in NB to an OD = 0.1, diluted 1/10 into DMEM + 10% FBS to opsonize the bacteria, and allowed to incubate statically for 1 h at 37⁰C and 5% CO_2_. Bacteria were then further diluted to achieve a multiplicity of infection (MOI) of 10 bacteria cells per macrophage in each well (in 100 μL) for the STm SGD mutant experiment or an MOI of 15 in the M9I-adapted STm strain experiment (see below for extra details). The plates were centrifuged at 500 × g for 2 min to sediment the STm cells and increase contact with the macrophages, before incubating at 37⁰C and 5% CO_2_. This was designated time zero. After 30 min of infection, the media was removed and replaced with fresh DMEM (10% FBS) containing 100 μg/mL gentamicin to kill the extracellular bacteria. The plates were incubated for 30 min at 37⁰C in 5% CO_2_, before removing and replacing the media once again with DMEM (10% FBS) with 10 μg/mL gentamicin. The cells were incubated as before for ∼20 h. To count the amount of intracellular STm, the macrophages were first washed with phosphate-buffered saline (PBS; 2 × 200 µL per well) and then lysed with 0.1% Triton X-100 in PBS, scraping using a pipette tip and forceful pipetting without the formation of bubbles. The lysates were diluted as needed in PBS and plated on NA for CFU counting. To minimize random error, the experiment was performed in two separate and identical plates, the samples of which were combined before plating. Within each plate, 3 biological replicates (from different initial bacterial cultures) were included in technical duplicate, and the experiment was repeated three or four times (n = 18-24). Outliers were tested and removed using the ROUT method on Graphpad Prism and statistical differences were calculated using a one-way ANOVA with multiple comparisons (for the STm SGD mutants) or a two-tailed t-test (for the M9I-adapted mutant). Due to high variability in the MOI across experiments (from the random error associated with diluting bacterial suspensions), in parallel with macrophage infection, the bacteria used for the infection were enumerated by serially diluting in PBS and plating on NA for CFU counting to confirm the true MOI of the sample. Outliers were tested using the ROUT method and infection results were removed from samples with outlier MOIs. A one-way ANOVA or two-tailed t-test were used to ensure no significant difference (p > 0.989) between conditions (**S5 Fig**).

### Whole Genome Sequencing of M9A- and M9I-adapted STm

Growths started from a single colony on NA, which was allowed to grow to stationary phase in NB prior to being subcultured 1/100 in either M9A or M9I, each in triplicate. A sample of the starting culture in NB was taken and the DNA was extracted using a Bio Basic (Canada) EZ-10 genomic DNA miniprep kit. M9A/M9I-adapted cultures were grown until they reached stationary phase before subculturing. After three subcultures, cells were collected, and their DNA was extracted for sequencing. Whole genome sequencing was performed at SeqCenter (Pittsburgh, PA, USA) using the Illumina NovaSeq X Plus. The sequencing depth was on average 350×. The Breseq v0.37.1 pipeline [109] was used for variant calling with the polymorphism flag. The genomes were deposited on the NCBI database, Submission ID (SUB15249457), BioProject ID (PRJNA1249104). The reference sequence for *Salmonella enterica* serovar Typhimurium ATCC 14028 used for variant calling is available at: genomes.atcc.org<u>/genomes/814482244337467e?tab=overview-tab</u>.

### STm growth and extraction for metabolomics

Media-adapted cultures (as described above) were washed once with their respective media before resuspending in 50 mL cultures in 250 mL Erlenmeyer flasks at an OD of 0.001 for MHB, or 0.1 for M9A and M9I before allowing to grow at 37⁰C, 250 rpm. The OD at stationary phase for each media under these conditions was established as 2.5 for MHB, 1.6 for M9A, and 1.1 for M9I. Samples were taken at two phases of growth: mid-exponential phase (EP) samples were collected when cultures were at 0.3× the stationary OD, and early-stationary phase (SP) samples were collected when cultures were at 0.9× the stationary OD for the respective media. Once cultures reach the appropriate ODs, a sample was taken and diluted in the appropriate media to get 13 mL of OD = 0.25, which allowed us to normalize how much biomass was collected across all conditions. The cells were pelleted by centrifugation in a 15 mL tube at 14 000 × g and 4⁰C for 10 min. All but 1.5 mL of the supernatant was removed, then the cells were resuspended and transferred to a 2 mL microfuge tube to be pelleted again. The cells were washed 1× with ice-cold 150 mM ammonium formate, pH 7.4, before centrifugation at 21 000 × g and 4⁰C for 5 min, before removing the supernatant. The cells were frozen at -80 ⁰C for 1-2 days before extraction. Additionally, no-cell media controls were made by processing a sample of each media the same as cellular pellets to identify background ion signals present in the media rather than the cell extracts.

The samples were always kept on ice during the extraction process. Cell pellets were lysed and extracted by first resuspending them in 380 μL of 50% MeOH/H_2_O (v/v), before adding 220 µL of MeCN and 6 ceramic beads (1.4 mm) per sample. Cells were lysed and homogenized by bead-beating for 2 minutes at 30 Hz using a QIAGEN TissueLyser II (Germany). Next, 600 μL of ice-cold DCM and 400 μL of ice-cold water were added to each sample, and the mixture was vortexed for 1 min before allowing the metabolites to partition on ice for 10 min. Lastly, the mixtures were centrifuged at 1500 × *g* and 1⁰C for 10 min. A 700 μL aliquot of the aqueous layer was collected and dried by vacuum centrifugation (Labconco, USA) at -4°C before being stored at -80 ⁰C until analysis.

### Data collection for targeted metabolomics of short chain CoA metabolites

For analysis of short chain CoA metabolites, samples were injected onto an Agilent 6470 Triple Quadrupole (QQQ)–LC–MS/MS (Agilent Technologies). Chromatographic separation of the metabolites was achieved using a 1290 Infinity ultra-performance quaternary pump liquid chromatography system (Agilent Technologies, USA). The mass spectrometer was equipped with a Jet StreamTM electrospray ionization source, and samples were analyzed in negative mode. The source-gas temperature and flow were set at 300°C and 5 L/min, respectively, the nebulizer pressure was set at 45 psi, and the capillary voltage was set at 3,500 V. Multiple reaction monitoring parameters (qualifier/quantifier ions and retention times) were optimized and validated using authentic metabolite standards. Metabolite standards were run at multiple stages along the data acquisition workflow, which were used as quality control (QC) samples in the analysis.

Dried extracts were re-suspended in 50 μL of chilled H_2_O, clarified by centrifugation at 1°C, and transferred to glass inserts before injection. Sample injection volumes for analyses were 5 μL. Chromatographic separation was achieved using a XBridge C18 column 3.5 μm, 2.1 × 150 mm (Waters). The chromatographic gradient started at 100% mobile phase A (10 mM ammonium acetate in water) with a 15 min gradient to 35% B (MeCN) at a flow rate of 0.2 mL/min. This was followed by an increase to 100% B over 0.1 min, a 4.9 min hold time at 100% mobile phase B and a subsequent re-equilibration time (6 min) at 100% A before the next injection.

### Data collection for ion pairing targeted metabolomic of central carbon metabolism

After extracts were sampled for CoA metabolites, they were analyzed using an ion pairing method on the same instrument as described above. The source-gas temperature and flow were set at 150°C and 13 L min^−1^, respectively, the nebulizer pressure was set at 45 psi, and the capillary voltage was set at 2,000 V. As above, QC samples made up of authentic metabolite standards were used to validate parameters.

Chromatographic separation of the isomers and other metabolites was achieved by using a Zorbax Extend C18 column 1.8 μm, 2.1 × 150 mm with guard column 1.8 μm, 2.1 × 5 mm (Agilent Technologies). The chromatographic gradient started at 100% mobile phase C (97% water, 3% methanol, 10 mM tributylamine, 15 mM acetic acid, 5 µM medronic acid) at a flow rate of 0.25 mL min^−1^ for 2.5 min, followed by a 5-min gradient to 20% mobile phase D (methanol, 10 mM tributylamine, 15 mM acetic acid, 5 µM medronic acid), a 5.5-min gradient to 45% D,a 7-min gradient to 99% D, and finally, a 4-min hold at 100% mobile phase D. The column was restored by washing with 99% mobile phase E (90% aqueous MeCN) for 3 min at 0.25 mL min^−1^, followed by increasing the flow rate to 0.8 mL min^−1^ over 0.5 min and holding for 3.85 min, before returning to 0.6 mL min^−1^ over 0.15 min. The column was then re-equilibrated at 100% C over 0.75 min, during which the flow rate was decreased to 0.4 mL min^−1^, and held for 7.65 min. One minute before the next injection, the flow rate was brought back to 0.25 mL min^−1^. The column temperature was maintained at 35°C. The metabolite standards used for metabolomics were purchased from Sigma-Aldrich.

### Metabolomics Analysis

Peak area data was carefully reviewed to ensure the accuracy and robustness of the analysis. Of the 296 metabolites in the targeted screen panel, ions that had weak or no signal were removed before analysis. The injection blanks and the controls made up of extracts from growth media alone were used to identify and remove metabolite signals that could not be resolved from the background.

Metaboanalyst 6.0 [110] was used to perform all metabolomics analyses unless otherwise stated. The data was filtered for reliability and to remove non-informative variables by excluding features that exhibited relative standard deviations (RSDs) greater than 30% of the QC samples, metabolites outside of the 5% interquantile range across all samples, and mean intensities within 5% of the limit of detection. The data was normalized to ensure a Gaussian distribution before analysis by log_10_ transforming and auto scaling the data. Multiple analyses were performed to ensure the robustness of the interpreted results by globally fitting the six experimental conditions in addition to analyzing important paired conditions. In the multiple variable analyses (across all six conditions), significant metabolites were identified by two-way ANOVA (p < 0.05). In the pairwise comparisons, T-test (p < 0.05) and fold-change (log_10_(FC) > 2) were used. Volcano plots and heatmaps were made using Graphpad prism and the calculated results or normalized abundances were computed using Metaboanalyst.

## Supporting Information

**S1 Fig**: **Growth data of various STm SPI-13 mutants in M9A and M9I.** This data was summarized in Figure 1b. Error bars depict the SD. n =3.

**S2 Fig**: **Gentamicin protection assay comparing the infection of STm wt and Δ*ich* from 1 h to 72 h.** Dotted lines and coloured bands depict the SD around each point connected by a straight line between point. n=4 per timepoint.

**S3 Fig: Overlap of genetic mutations within the three replicates of STm adapted to (A) M9A or (B) M9I.** Genes with mutations that were found at frequencies below 25% are coloured in red and are not bolded, while those found at frequencies above 25% that may have conferred an advantage for growth in that specific media are in bold.

**S4 Fig: Comparing relevant pairs of conditions in the metabolomic experiments.** (A) Volcano plots for pairwise comparisons between M9I vs M9A or MHB and M9A vs MHB at both EP and SP. Highlighted metabolites were considered significant when log10(FC) > 2 and p-value < 0.05. (B) Venn diagram showing the overlap of significant metabolites between the three media (after combing the EP and SP results for each media). Results were stratified to better understand the effects on different pathways by pooling significant features from different comparisons and performing a pathway enrichment analysis. (C) “M9 vs MHB” pooled significant features in the M9A/I vs MHB (EP and SP) comparisons and (D) “M9I vs other” pooled both sets of M9I vs MHB/M9A (EP and SP) comparisons. **S5 Fig**: **MOI of the inoculum used for gentamicin protection assay.** (A) Comparing STm SPI-13 mutants to wt STm and (B) comparing STm wt to M9I-adapted STm.

S1 Table: Background mutations observed prior to the M9A/M9I-adaptation experiments after initial growth in MHB.

S1 Appendix. Compilation of all supporting figures and table referenced in the main text of this study. Includes captioned images of S1 Fig, S2 Fig, S3 Fig, S4 Fig, S5 Fig, S1 Table.

S1 Data. Raw peak intensity data used for the metabolomics analysis presented in this study.

## Acknowledgements

We would like to thank Prof. Eric Brown at McMaster University for gifting us the *Salmonella* Typhimurium mutant strains. Metabolites were analysed at the Rosalind and Morris Goodman Cancer Institute’s Metabolomics Innovation Resource, McGill University. This facility is supported by the Terry Fox Foundation, Quebec Breast Cancer Foundation, The Dr. R. John Fraser and Mrs. Clara M. Fraser Memorial Trust, and McGill University.

## Author Contributions

**Conceptualization**: Jacob Pierscianowski, Karine Auclair

**Data curation**: Jacob Pierscianowski, Farhan R. Chowdhury

**Formal analysis**: Jacob Pierscianowski, Farhan R. Chowdhury

**Funding acquisition**: Andréanne Lupien, Brandon L. Findlay, Karine Auclair

**Investigation**: Jacob Pierscianowski, Suzana Diaconescu-Grabari, Farhan R. Chowdhury

**Methodology**: Jacob Pierscianowski, Farhan R. Chowdhury, Brandon L. Findlay, Karine Auclair **Project administration**: Karine Auclair

**Resources**: Jacob Pierscianowski, Suzana Diaconescu-Grabari, Andréanne Lupien, Karine Auclair

**Supervision**: Brandon L. Findlay, Karine Auclair

**Validation**: Jacob Pierscianowski, Karine Auclair

**Visualization**: Jacob Pierscianowski

**Writing –original draft**: Jacob Pierscianowski

**Writing –review & editing**: Jacob Pierscianowski, Farhan R. Chowdhury, Brandon L. Findlay, Karine Auclair

## Data Availability Statement

All relevant data are within the paper and its Supporting Information.

## Funding

This research was funded by the Canadian Institute of Health Research (CIHR grant PJT-166175) to K.A. and the FRQ-funded Réseau Québecois de Recherche sur les Médicaments (RQRM grant 78855) to K.A. and B.L.F. NSERC and FRQS are also thanked for scholarships to J.P. (CGSM and PGSD) and F.R.C. (B2X Graduate Studentship), respectively. The funders had no role in study design, data collection and analysis, decision to publish, or preparation of the manuscript.

## Competing Interests

The authors declare no competing interests.

## Notes

### Competing Interest Statement

The authors have declared no competing interest.

### Summary of Updates

The Manuscript was submitted by the Journal without the Supporting Information. I am now adding the Supporting Information.

