## Supporting Information for "Metabolic and genomic adaptations of *Salmonella* Typhimurium grown on itaconate"

**S1 Appendix. Compilation of all supporting figures and tables referenced in the main text of this study.**

### Table of Contents

S1 Fig: Growth data of various STm SPI-13 mutants in M9A and M9I.

S2 Fig: Gentamicin protection assay comparing the infection of STm wt and Δ*ich* from 1 h to 72 h.

S3 Fig: Overlap of genetic mutations within the three replicates of STm adapted to (A) M9A or (B) M9I.

S4 Fig: Comparing relevant pairs of conditions in the metabolomic experiments.

S5 Fig: MOI of the inoculum used for gentamicin protection assay.

S1 Table: Background mutations observed prior to the M9A/M9I-adaptation experiments after initial growth in MHB.

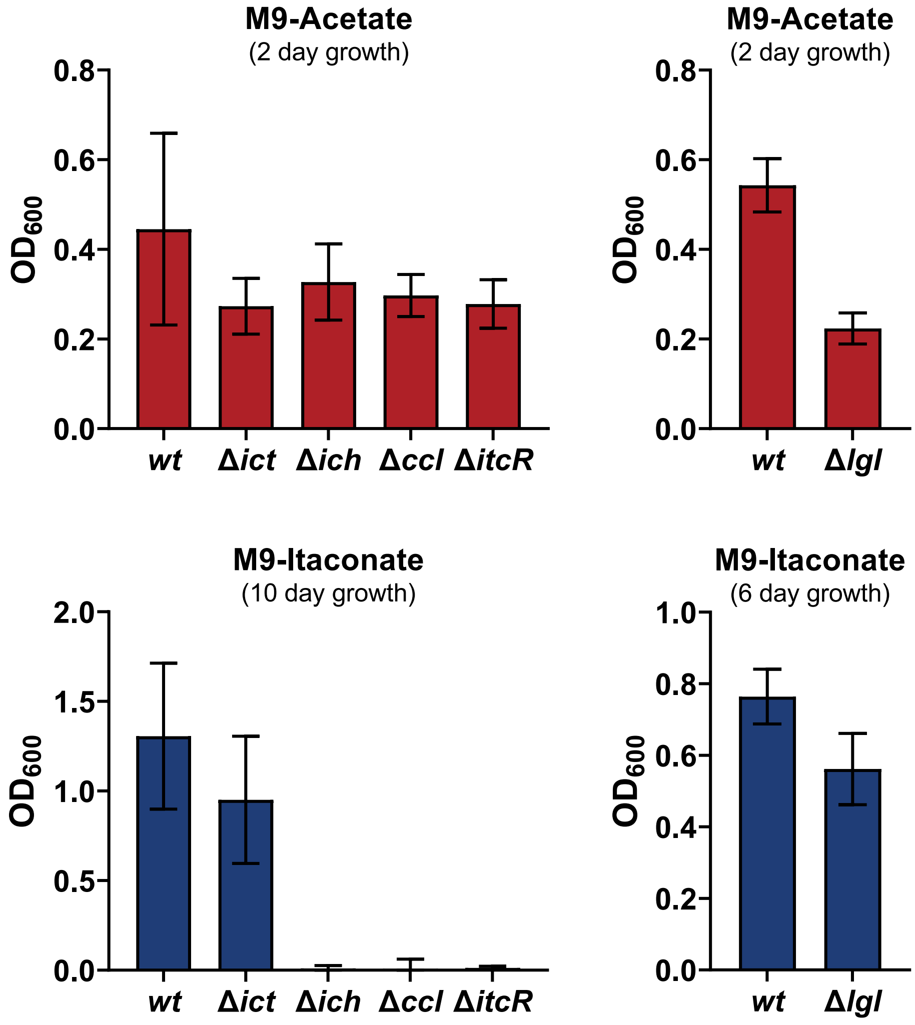


**S1 Fig**: **Growth data of various STm IRO mutants in M9A and M9I.** This data was summarized in Figure 1b. Error bars depict the SD for n = 3.

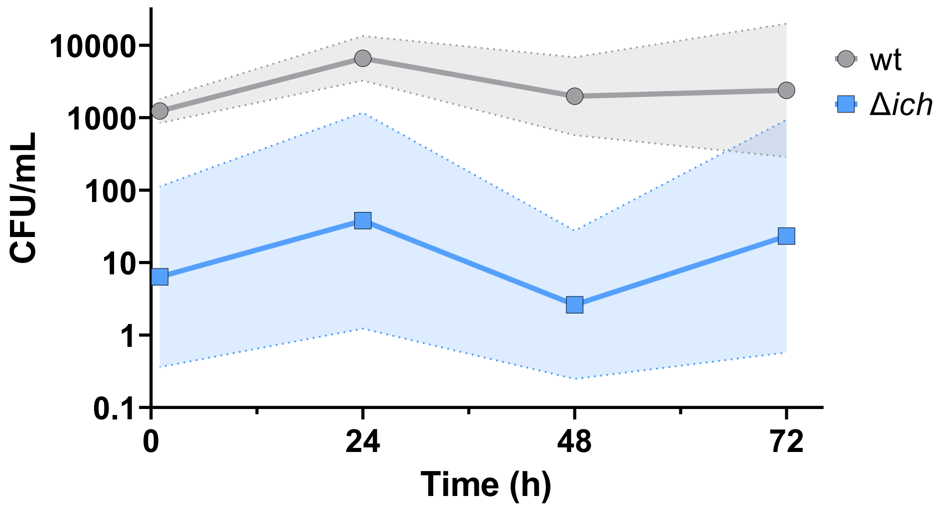


**S2 Fig**: **Gentamicin protection assay comparing the infection of STm wt and Δ*ich* over 72 h.** The coloured bands depict the SD around each datapoint, which are connected by a straight line; n = 4 per timepoint.

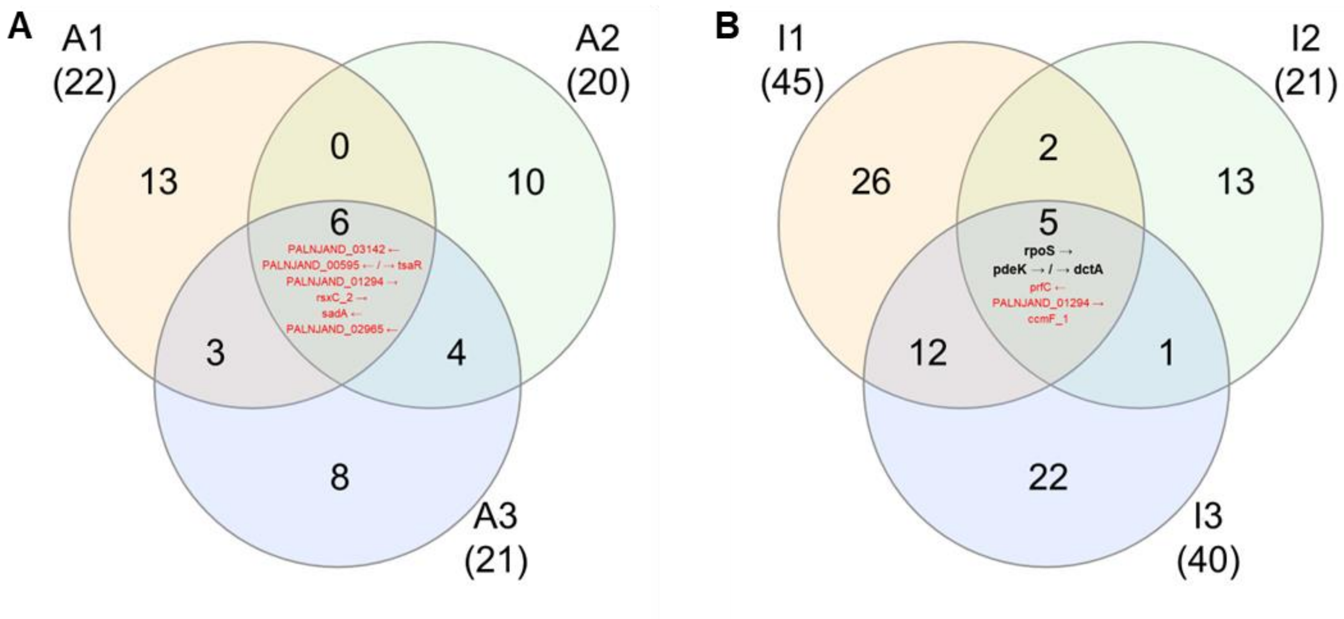


**S3 Fig: Overlap of genetic mutations within the three replicates of STm adapted to (A) M9A or (B) M9I.** Genes with mutations that were found at frequencies below 25% were are coloured in red and are not bolded, while those found at frequencies above 25% that may have conferred an advantage for growth in that specific media are in bold.

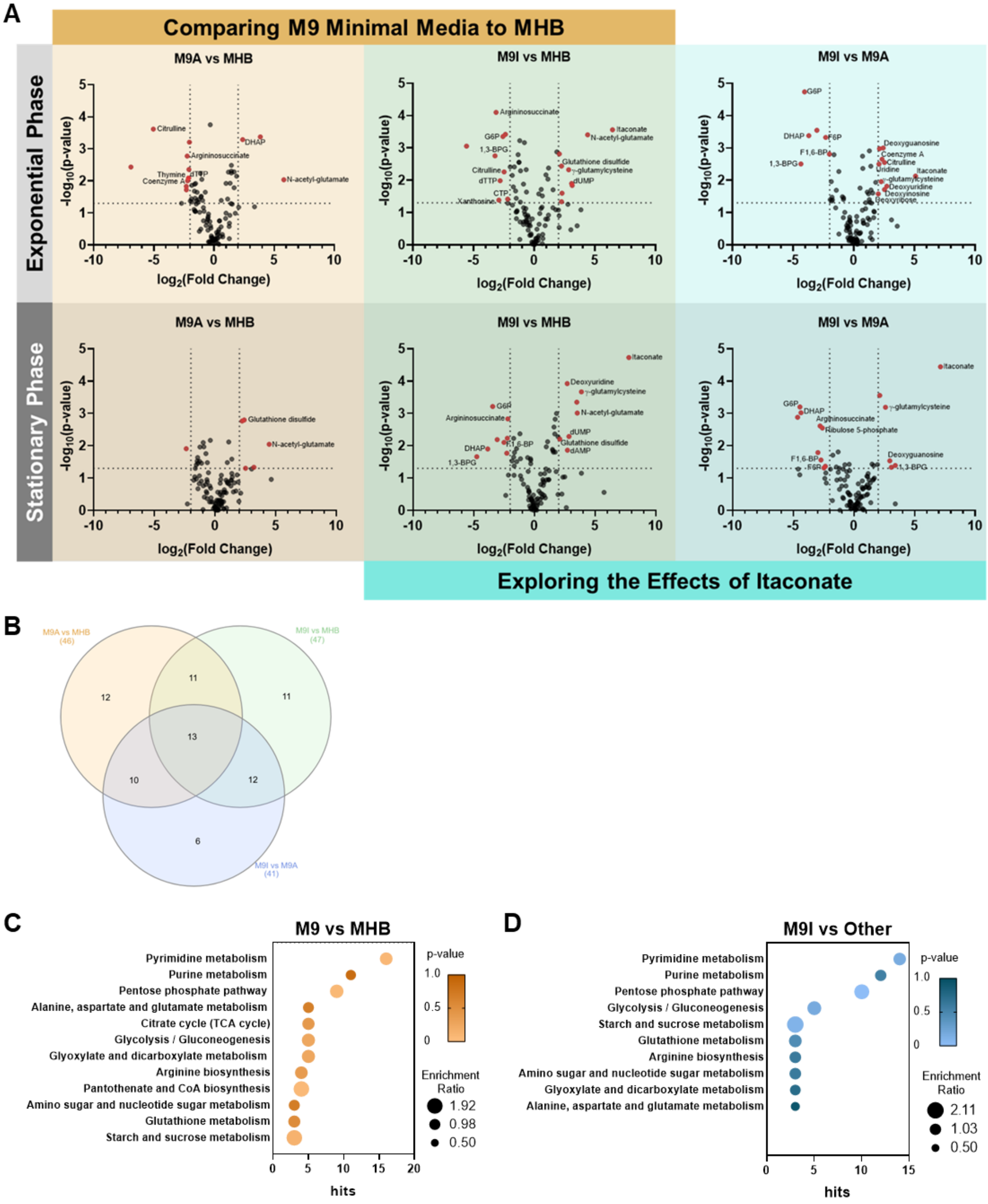


**S4 Fig: Comparing relevant pairs of conditions in the metabolomic experiments.** (A) Volcano plots for pairwise comparisons between M9I vs M9A, or MHB and M9A vs MHB at both EP and SP.

Highlighted metabolites were considered significant when $\log_{10}(FC) > 2$ and p-value < 0.05. (B) Venn diagram showing the overlap of significant metabolites between the three media (after combing the EP and SP results for each media). Results were stratified to better understand the effects on different pathways by pooling significant features from different comparisons and performing a pathway enrichment analysis. (C) "M9 vs MHB" pooled significant features in the M9A/I vs MHB (EP and SP) comparisons and (D) "M9I vs other" pooled both sets of M9I vs MHB/M9A (EP and SP) comparisons.

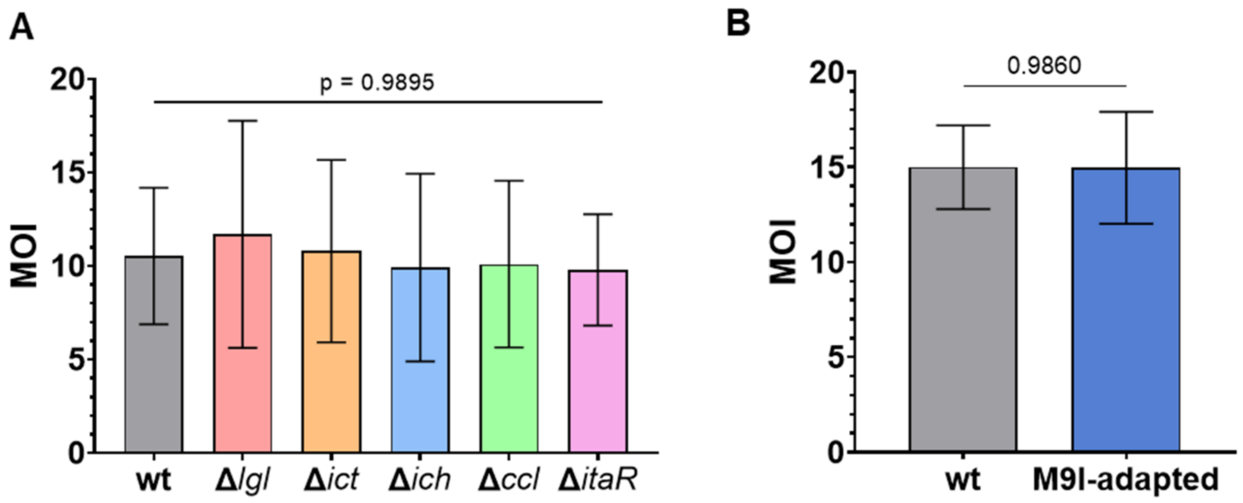


**S5 Fig**: **MOI of the inoculum used for gentamicin protection assay.** (A) Comparing STm IRO mutants to wt STm and (B) comparing wt STm to M9I-adapted STm.

**S1 Table**: **Background mutations observed prior to the M9A/M9I-adaptation experiments after initial growth in MHB.**

| Gene | Mutation | Type | Frequency | Description |
|---|---|---|---|---|
| ***nadB*** **←** | I530F (ATC→TTC) | Missense | 100% | L-aspartate oxidase |
| ***sfmH_1*** **←** | I157F (ATT→TTT) | Missense | 100% | putative fimbrial-like protein SfmH |
| **PALNJAND_02872 ←** | S1002N (AGC→AAC) | Missense | 45% | hypothetical protein |
| **PALNJAND_00365 ←** | P202P (CCT→CCC) | Silent | 16% | hypothetical protein |
| | G175D (GGC→GAC) | Missense | 11% | |
| | D164D (GAT→GAC) | Silent | 10% | |
| | D225D (GAT→GAC) | Silent | 9% | |
| | P147P (CCT→CCC) | Silent | 8% | |
| | D236D (GAT→GAC) | Silent | 7% | |
| | V199V (GTG→GTA) | Silent | 6% | |
| | S139S (AGT→AGC) | Silent | 5% | |
| ***rsxC_2*** **→** | A707V (GCC→GTC) | Silent | 13% | Electron transport complex subunit RsxC |
| | A688A (GCT→GCC) | Silent | 11% | |
| | E684E (GAA→GAG) | Silent | 9% | |
| | D708D (GAC→GAT) | Silent | 7% | |
| | A685A (GCG→GCA) | Silent | 7% | |
| | E622E (GAA→GAG) | Silent | 6% | |
| | P706A (CCG→GCG) | Missense | 6% | |
| ***sadA*** **←** | T549T (ACA→ACC) | Silent | 11% | Autotransporter adhesin SadA |
| | S603N (AGT→AAT) | Missense | 9% | |
| | L620L (TTA→TTG) | Silent | 7% | |
| | A545D (GCC→GAC) | Missense | 7% | |
| | A467A (GCT→GCG) | Silent | 7% | |
| | K547K (AAG→AAA) | Silent | 7% | |
| | A527A (GCC→GCA) | Silent | 7% | |
| | N540K (AAC→AAG) | Missense | 5% | |
| **PALNJAND_03142 ←** | A241E (GCG→GAG) | Missense | 13% | hypothetical protein |
| | E209D (GAA→GAC) | Missense | 9% | |
| | E164D (GAA→GAC) | Missense | 9% | |
| | K146K (AAA→AAG) | Missense | 8% | |
| | V168A (GTA→GCA) | Missense | 7% | |
| | A171A (GCA→GCG) | Silent | 6% | |
| | E149D (GAG→GAT) | Missense | 5% | |
| **PALNJAND_01294 →** | G929G (GGT→GGC) | Silent | 11% | hypothetical protein |
| | T714I (ACC→ATC) | Missense | 8% | |
| | D743E (GAT→GAG) | Missense | 7% | |
| | G798G (GGC→GGT) | Silent | 6% | |
| | G767D (GGC→GAC) | Missense | 5% | |
| | V802V (GTC→GTG) | Silent | 5% | |
| **PALNJAND_00595 ← / →** ***tsaR*** | intergenic (-328/-892) (G→A) | Missense | 12% | hypothetical protein/HTH-type transcriptional regulator TsaR |
| | intergenic (-212/-1008) (G→A) | Missense | 9% | |
| | intergenic (-431/-789) (C→T) | Missense | 9% | |
| ***prfC*** **←** | A278T (GCG→ACG) | Missense | 17% | Peptide chain release factor RF3 |
| | R287H (CGC→CAC) | Missense | 12% | |
| **PALNJAND_02099 → / ← PALNJAND_02100** | intergenic (+408/+376) (G→A) | Missense | 5% | hypothetical protein/tRNA-Tyr |
| | intergenic (+503/+281) (G→T) | Missense | 5% | |
